# Using Photoswitching FRET to Define the Interaction Boundaries between the Rab1b GTPase and Secretory Cargo

**DOI:** 10.1101/2024.06.23.600248

**Authors:** Yehonathan Malis, Guy M. Hirschberg, George H. Patterson, Koret Hirschberg

## Abstract

FRET is a powerful tool to simultaneously establish and localize interactions between fluorescently tagged proteins with high spatial resolution. Rainey K.H. and Patterson G.H. introduced Photoswitching FRET (psFRET) using the Dronpa Photoswitching fluorescent protein. We present a straightforward detailed method, and a powerful software tool that allows adaptation of psFRET to diverse experimental setups. Image stacks, recording the decay of the Dronpa donor, serve as input to the software utility that includes effective preprocessing options preceding the calculation FRET efficiency at the single pixel level. We applied psFRET to generate interaction maps analyzing diverse interactions between cargo proteins, the GTPase Rab1b, and GRASP65 during ER to Golgi trafficking. Cargo-Rab1b interactions were restricted to the transit period from ER to Golgi. These data lend support to a mechanism whereby cargo sensing may regulate the level of downstream effectors recruitment to secretory membranes by Rab1.

## Introduction

For biologists, FRET is a powerful tool to establish and localize interactions between fluorescently tagged proteins with high spatial resolution. Numerous methods to detect FRET with varying reliability and ease of use exist. For example, fluorescence lifetime imaging that requires costly dedicated equipment and complex analysis on one end, or acceptor photobleaching that is relatively straightforward but depends on the donor and acceptor molecules’ fluorescence intensity on the other. Photo-switching FRET (psFRET), using the Dronpa Photo-switching fluorescent protein, was recently introduced by Rainey K.H. and Patterson G. H. (Rainey and Patterson, 2019). In this study, we formulated a detailed experimental protocol. Furthermore, we developed an application that will carry out the accompanying extensive calculations required, thereby rendering psFRET usable by the large readership of cell biologists. To this end, we applied this method to study interactions among transport machinery proteins and secretory cargo at the early stages of anterograde transport. The psFRET is based on the decrease in the decay rate of Dronpa fluorescent protein that results from energy transfer to an acceptor molecule, a red fluorescent protein. The decay of Dronpa is not photobleaching but rather a reversible switch-off that can be reversed by exposing Dronpa to a 405 nm laser. Thus, experiments can be repeated multiple times, and fast imaging equipment may allow analysis FRET dynamics in living cells. In brief, a series of images of cells expressing Dronpa and mCherry-tagged potential interacting proteins are captured using enough laser intensity to gradually allow the Dronpa donor to completely switch off. The acceptor fluorescence in a small ROI within the image is photobleached to obtain the Dronpa decay coefficient values without an acceptor. These image stacks serve as the input to the software we designed that can generate the Dronpa exponential decay coefficient by fitting it to an exponential equation at the single-pixel level. The software includes various traditional powerful image preprocessing capabilities. The output also consists of a detailed statistical evaluation of the decay coefficients obtained. This output is straightforwardly translated to a FRET efficiency pseudo-color interaction map. To test the potential of the psFRET, we asked to primarily study interactions among transport machinery and cargo proteins during ER to Golgi transport. We picked Rab1b as it is a well-characterized small GTPase associated with regulating ER export, and its GTP-bound form was demonstrated to colocalize with cargo proteins on early secretory membranes, including ER-exit sites (ERESs), transport carriers, ERGIC, and Golgi membranes. Rab1b is a small GTPase from the Rab GTPases protein family. Its exchange of GDP to GTP and hydrolysis are mediated by a set of GDP exchange factors (GEFs) and GTPase activating proteins (GAPs). Rab proteins have a lipid modification that, upon GTP binding, is exposed and promotes membrane binding. The membrane-associated GTP-bound form of Rab1b is known to recruit numerous downstream effectors, such as the tethering and docking proteins GRASP65, p115, GM130, and Giantin (Beard et al., 2005; Moyer et al., 2001; Rosing et al., 2007). As Rab1b-GTP binds to ERES, ER to Golgi carriers and Golgi membranes (Westrate et al., 2020), and as it was shown to be involved in COPI coat dynamics via recruiting GBF1, the GEF for ARF1 GTPase, Rab1b was identified as a master regulator that can affect the rate of bulk cargo export from the ER. However, the mechanism underlying this function needs to be clarified.

Here, we used psFRET to study the interaction of Rab1b with its downstream effector GRASP65 and the cargo protein vesicular stomatitis virus G protein (VSVG). Rab1b-VSVG interaction was limited to the early ER to Golgi trafficking stage. Based on the psFRET in the Golgi, this interaction ceased regardless of the high donor and acceptor concentrations present.

## Results

The psFRET method using the Dronpa photo-switchable fluorescent protein was recently introduced. The basis for the technique is that the decay rate of the Dronpa protein is reduced in the presence of an effective FRET acceptor, such as any red fluorescent protein. The Dronpa decay is not photobleaching but rather a reversible photoconversion, and thus the protein can be switched on repeatedly using, for example, a 405 nm laser line. Here, we developed an experimental protocol fit for a wide range of potential users, enabling application of psFRET to create a FRET interaction map at pixel level resolution. A detailed experimental protocol is provided in the methods section and illustrated in Fig. 1. After cells are fixed, the field of view to be analyzed is chosen so that it will contain an area within which the acceptor fluorescence can be photobleached to obtain the Dronpa decay coefficient in the absence of FRET (*K_D_*, no FRET acceptor molecule). This Dronpa decay coefficient in the absence of FRET is required for calculating FRET efficiency. A series of approximately 10 images is captured to record the stepwise decay of the Dronpa donor. Fig. 2 displays the use of the experimental protocol and the analysis using the Dronpa-5-mCherry, a Dronpa donor, and a mCherry acceptor linked by five amino acids (Rainey and Patterson, 2019). Fig. 2A shows the field of view that includes the photobleached region. Images are then captured with standard laser power, causing the gradual photoconversion of the Dronpa to its dark state. The experiment can be repeated multiple times by switching on the Dronpa protein using a 405 nm laser line (Figure 2B). Each stack is then analyzed by the software utility developed to fit the Dronpa decay to an exponential equation. The fitting process is carried out on the entire image to extract the exponential decay coefficient for each individual pixel. The psFRET utility software has several preprocessing options that can be performed using pooled or sliding kernels with varying sizes (Fig. S1A). The output of the psFRET utility includes four spreadsheets: The decay coefficients and three statistical parameters to evaluate the fit: Standard error, Chi-square, and Root Mean Squared Error (RMSE). The decay coefficients (*K_DA_*) for each pixel or kernelled pixels can be converted to FRET efficiency (FRET_eff_) using equation 1.

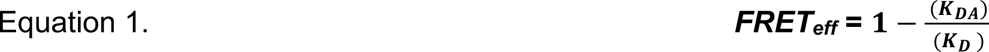

**Figure 1.**
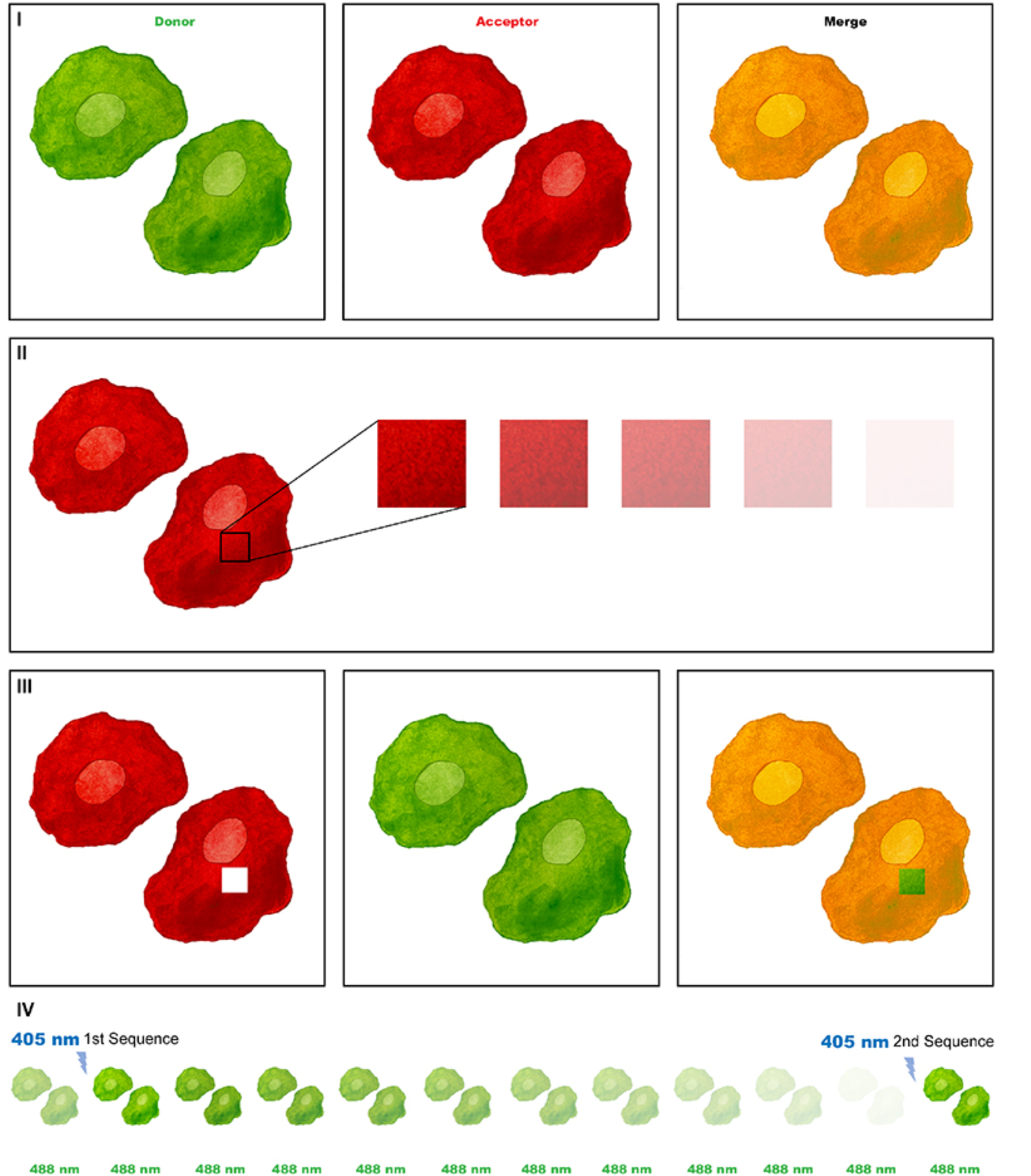
The psFRET experimental protocol. (I) Choose a field of view that contains two cells that express both donor and acceptor molecules. Capture image using settings that utilize the full dynamic range of the Dronpa donor channel. (II) To obtain the control decay coefficient in the absence of an acceptor, zoom into a high intensity square region within one of the cells and bleach the entire acceptor population. (III) Take a reference image of both green and red channels by reusing settings of panel I. (IV) Take a set of images using intensity that will gradually convert the Dronpa population to its dark state.

**Fig. 2.**
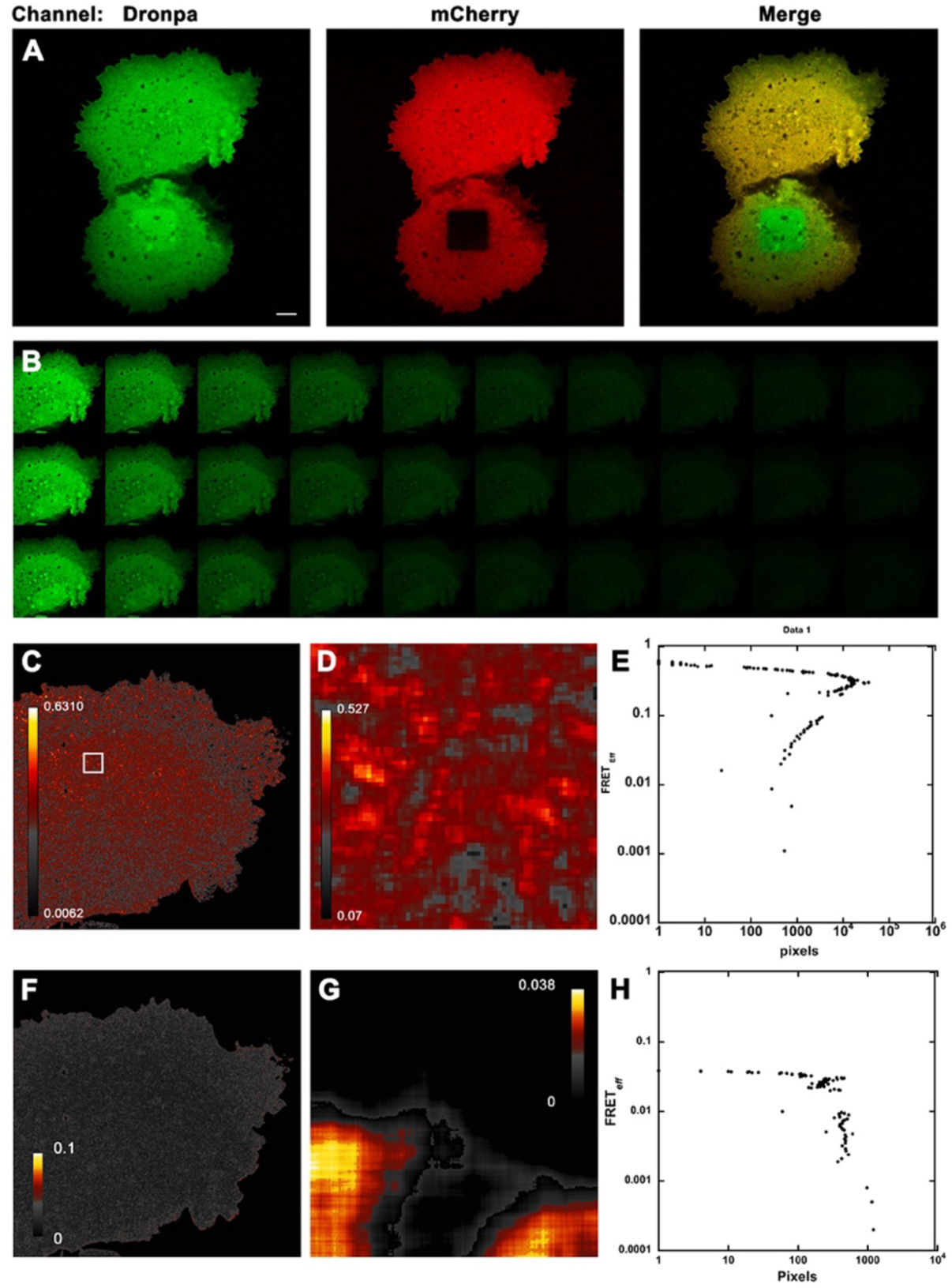
The psFRET method, psFRET analysis of the Dronpa-5-mCherry. **(A)** COS7 cells expressing the Dronpa-5-mCherry chimera were fixed using formaldehyde. The mCherry acceptor fluorescence was bleached using repetitive scanning with high power 561 nm laser line over a square ROI. **(B)** Three sets of input data in the form of image stacks each containing 10 images of the top cell in A captured after using 405 nm laser to switch-on the Dronpa protein. **(C)**. A pseudo color representation of FRETeff interaction map of the top cell in A, calculated using Equation 1. For this analysis the analyzed cell was processed using the uniform filter option of the software tool with a 2 x 2 kernel. The bleached box data was processed using the uniform filter option with an 8 x 8 kernel. Data worksheet was converted to a pseudo-color image using ImageJ as described in the “methods” section. **(D)** A 15-fold enlargement of the ROI in C. **(E)** A full log scale histogram showing the distribution of FRET efficiency values in the cell in C. **(F)** Same as C, a pseudo-color representation of the distribution of the standard error of the fit. **(G)** A pseudo-color representation of the distribution of FRET efficiency analyzed for the bleached box ROI. Lookup table shows FRETeff values between 0 % and 3.8 %. **(H)** Full log scale histogram of the data in G. Scale bars = 10 µm.

The spreadsheet containing the FRET_eff_ data is transformed to a pseudo color image to generate a full field interaction map using Image J (supplementary Fig. S1B) Figures 2C, 2D, and the full log scale histogram in 2E show the FRET_eff_ interaction map and the quantification of FRET_eff_ for the top cell in 2A using a kernel of 2 x 2 pixels, respectively. Figure 2F shows the standard error worksheet converted to a pseudo-color map. Finally, Figures 2G and 2H illustrate a FRET_eff_ interaction map and a full logscale FRET_eff_ histogram for the bleach box in the lower cell in 2A using an 8 x 8 kernel.

### psFRET analysis of Rab1b known interactors

We next used the psFRET to analyze the interaction between Rab1b and GRASP65, one of its known downstream effectors, a member of the Golgi tethering Rab1b effector complex, together with GM130 and P115 (Moyer et al., 2001). GRASP65 is a cytosolic protein that dynamically binds early secretory membranes. Three psFRET interaction maps are shown for three ROIs I through III (Fig. 3). Positive FRET is seen on all Rab1b-positive membranes, including the Golgi and peripheral tubular membranes (Dukhovny et al., 2008). The FRET occurs in distinct punctate structures. The positive FRET spots are roughly evenly distributed within Golgi membranes. (Fig. 3 II). The Rab1b positive membrane structure in Fig. 3 III resembles an ER to Golgi transport carrier based on size and organization and has the largest FRET puncta at its edge. It is tempting to speculate that these positive FRET zones may be associated with the tethering function of GRASP65.

**Fig. 3.**
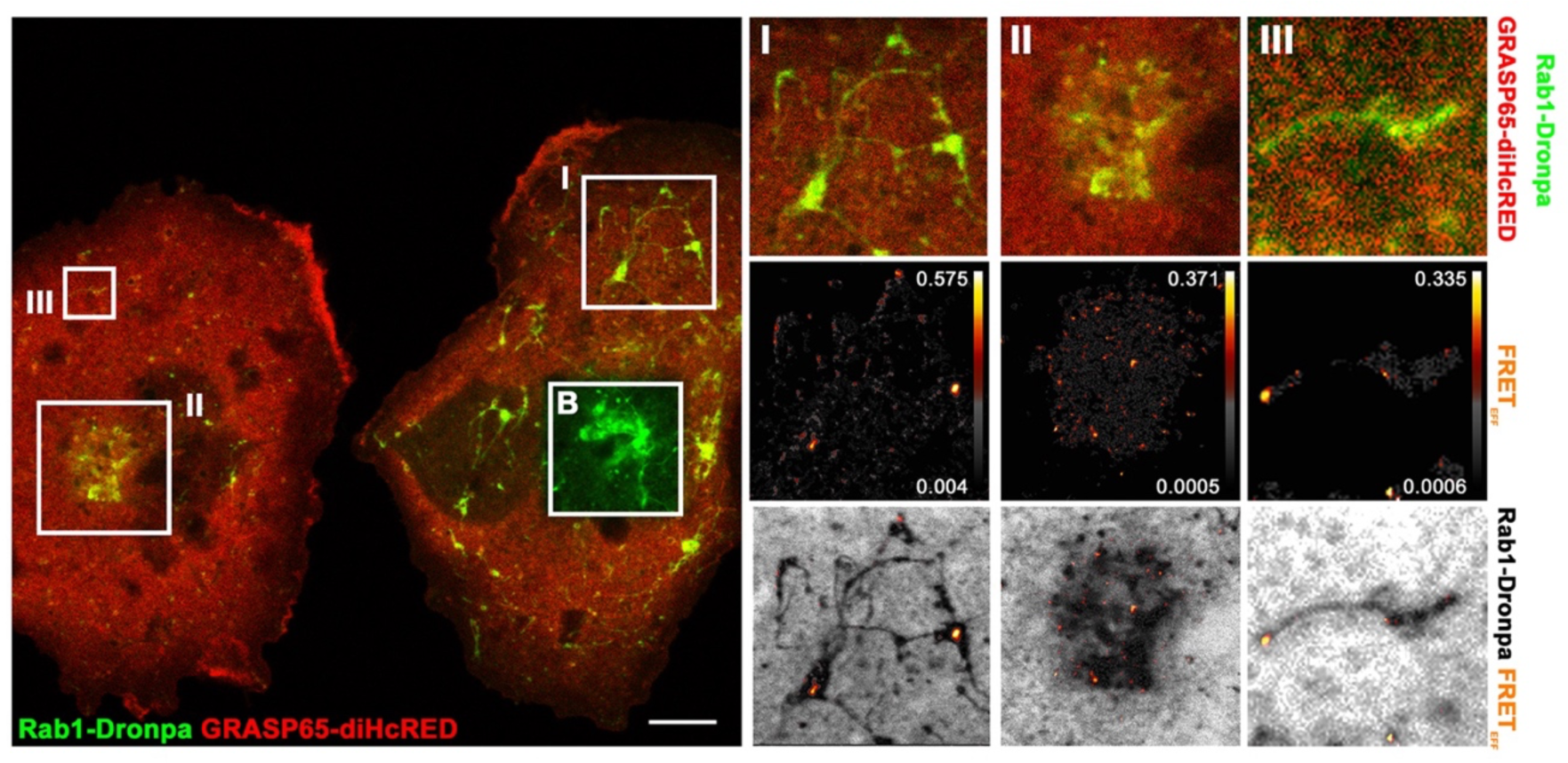
Interaction of Rab1b with its downstream effector GRASP65 using psFRET analysis. COS7 cells co-expressing Rab1b-Dronpa and GRASP65-diHcRED were fixed and analyzed as previously described. Confocal image shows two cells and four ROIs one of which is the bleach-box (B) over the Golgi apparatus of the cell at the right-hand side. ROIs I, II and III show a peripheral tubular structure, the Golgi apparatus, and a potential ER to Golgi transport carrier, respectively. Top panel is donor and acceptor fluorescent channels. Middle panel is the FRET efficiency interaction map and bottom panel is the interaction maps overlayed on the inverted donor channel. Lookup tables show range of FRET efficiency. Scale bar = 10 µm.

### Rab1b interacts with VSVG during ER to Golgi transport

Rab1b was demonstrated to colocalize with cargo-loaded membranes during ER to Golgi transport (Shomron et al., 2021; Westrate et al., 2020). Thus, we asked if the psFRET method may be applied to analyze potential Rab1b-Cargo interaction. Traditional acceptor photobleaching FRET analysis showed positive FRET between VSVG-GFP and Rab1b-mCherry. Here, we gradually photobleached the donor fluorescence within an ROI over the cell center, including the Golgi apparatus in a fixed cell coexpressing VSVG-GFP and Rab1b-mCherry. Before fixation, cells were transferred from non-permissive (39.5°C) to permissive (32°C) temperatures for about 30 min. Figure S2 shows a concurrence between donor fluorescence increase and acceptor photobleaching level, indicating FRET interaction. As opposed to psFRET where analysis is performed at the single pixel level, the analysis in Figure S2 was applied to the average fluorescence of the entire ROI.

We thus applied psFRET to utilize its superior spatial resolution. To this end, cells coexpressing the thermoreversible folding mutant of vesicular stomatitis virus G protein tagged with the scarlet red fluorescent protein (VSVG-Scarlet) and Rab1b-Dronpa were fixed after 0-, 20-, and 30-min incubations at the permissive temperature of 32°C following an overnight incubation at 39.5°C (Fig. 4). Before shifting to permissive temperature, VSVG is misfolded and is retained in the ER. At time points that include incubation at the permissive temperature of 32°C, it localizes to pre-Golgi (ER exit sites and transport carriers) and Golgi membranes (Hirschberg et al., 1998). Figure 4B shows the psFRET interaction maps of the cells in Fig. 4A for the 0- and 20-min after shift and the Golgi ROI (I) for the cell fixed at 30 min after shift. Fig. 4B demonstrates a substantial increase in Rab1b-VSVG psFRET efficiency after 20 min at the permissive temperature (32° C). Full log histograms analyzing psFRET efficiency show two orders of magnitude increase in the number of pixels with positive FRET efficiency and an increase in FRET efficiency values (Figure S3). Another distinct observation is that at 20 and 30 min after the shift, psFRET was distributed all over the cell as expected from the distribution of membrane structures that traffic from ER to Golgi but were essentially absent from the Golgi apparatus. During the 20- and 30-minute after the shift to permissive temperature time points, the fluorescence of both donor and acceptor is at high levels in Golgi membranes (Hirschberg et al., 1998). (Fig. 4B, images at the center and right-hand side). For comparison, an ROI (II) over part of the cell periphery showed psFRET interaction localized to peripheral tubular membranes (Fig. 4C). Thus, these data suggested that Rab1b-Cargo interaction is restricted to ER to Golgi transit time and is severed upon Golgi entry. The lack of FRET interaction in the Golgi demonstrates the independence of psFRET from fluorescence intensity as in the case of acceptor photobleaching.

**Fig. 4.**
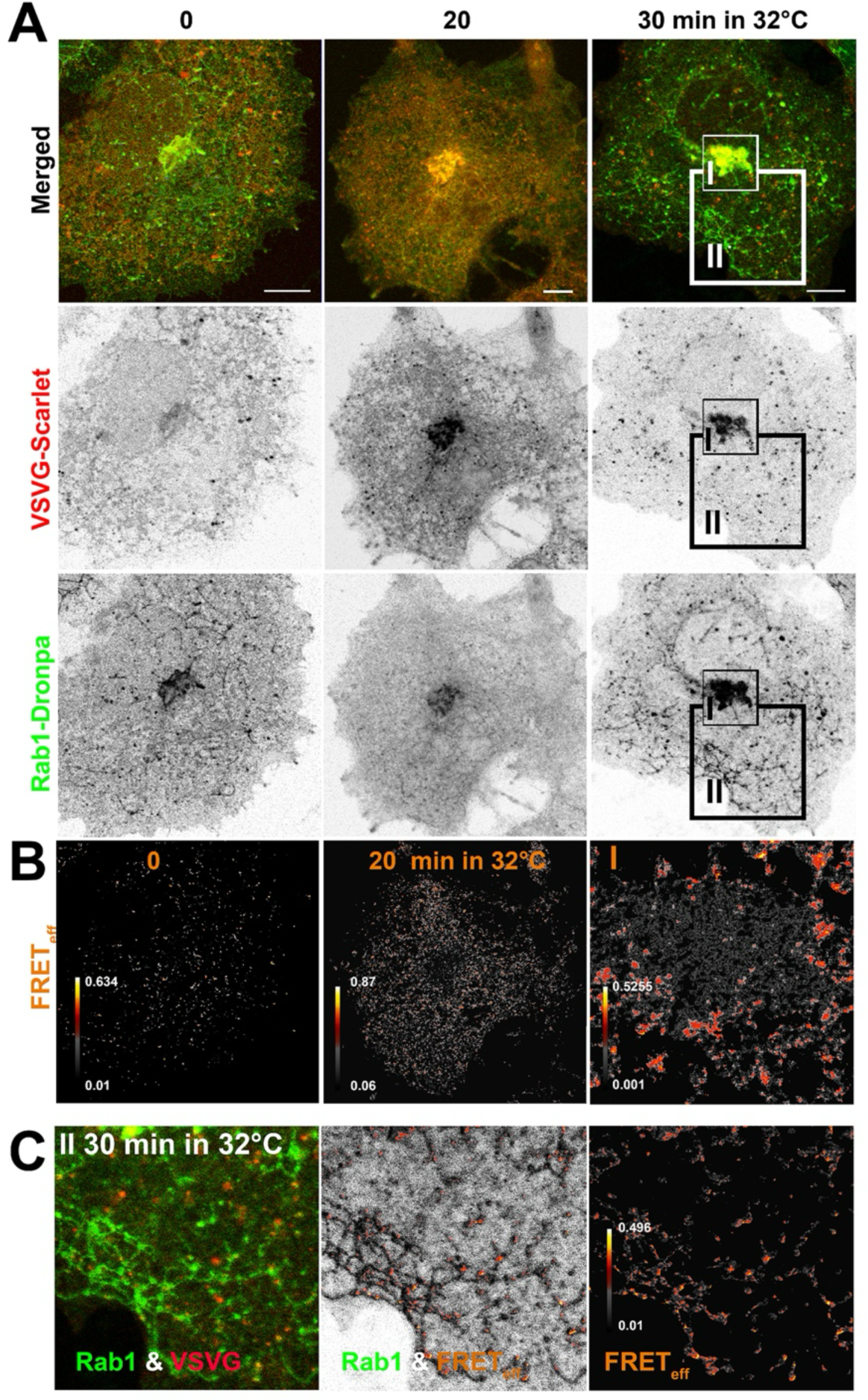
Rab1b-Dronpa and VSVG-Scarlet interact during ER to Golgi transport. **(A)** COS7 cells co-expressing Rab1b-Dronpa and VSVG-Scarlet were fixed after incubation for 0, 20, and 30 minutes at the permissive temperature of 32°C following overnight at the non-permissive temperature of 39.5 °C. Cells were fixed and analyzed as described. Top panel images are the merged donor (Rab1b-Dronpa, green) and acceptor (VSVG-Scarlet, red) fluorescence. ROI I and II at the right-hand side image are The Golgi and peripheral tubular membrane structure, respectively. Middle and bottom panels are inverted images of the acceptor and donor fluorescence, respectively. **(B)** FRET efficiency interaction maps for the cells incubated at 0 and 20 min at 32°C (left hand side and middle). Image at the right-hand side is a FRETeff interaction map of the Golgi ROI I. **(C)** Left to right, Donor and acceptor fluorescence, FRETeff over inverted donor image and FRETeff interaction map of ROI II. bars = 10 µm.

### Rab1b and VSVG do not interact in Golgi membranes

To further establish Rab1-VSVG interaction, we carried out a reciprocal experiment, switching the donor and acceptor tags. Figure 5A shows the entire field of view, including two cells fixed after 25 min at the permissive temperature (32°C) following an overnight incubation at the nonpermissive temperature (39.5°C). The top left cell contains the acceptor bleach box required to obtain the Dronpa decay control values. The Golgi ROI of the lower right-hand cell is analyzed for the donor (Fig. 5B) and acceptor (Fig. 5C) fluorescence, psFRET (Fig. 5D), and the standard error of the fit values (Fig. 6E). It is apparent that the psFRET between VSVG and Rab1b does not occur on Golgi membranes. In Figure 5F, the psFRET efficiency image is overlayed on top of the VSVG-Dronpa donor fluorescence channel. We analyzed five small ROIs over psFRET-positive pixels and a sixth one containing high donor and acceptor fluorescence values over the Golgi. We analyzed donor and acceptor fluorescence intensities ± SD, the psFRET efficiency ±SD, and the Standard error of the psFRET values ±SD (Fig. 5G) for each ROI. For example, ROI #1, with the area of 0.16µm^2^ (33 pixels), has mean donor and acceptor fluorescent intensities of 21 and 19 (8 bit) while average psFRET efficiency was 50.2 ± 16.6%. The average standard error of the fit for this ROI was 6.9 % ± 2.6%. For comparison, ROI #6 over the Golgi had mean donor and acceptor fluorescent intensities of 92 and 215 (8-bit), while psFRET efficiency was 10.9 ± 19.7%. The average standard error of the psFRET values for this ROI was 2.2 % ± 0.8%. These data confirm that there is little or no apparent FRET interaction between VSVG and Rab1b in Golgi membranes based on the psFRET method. Moreover, as FRET detects proximity it is unclear what is the contribution of crowding to the positive FRET signal. Here however, the evident lack of psFRET in Golgi membranes with high donor and acceptor fluorescent intensities dismisses the idea that the observed psFRET in pre-Golgi membranes indicates crowding rather than interaction (Malis et al., 2024). The Independence of FRET efficiency from acceptor and donor fluorescence eliminates the possibility of any contribution from the fluorescent tags to the apparent interaction.

**Fig. 5.**
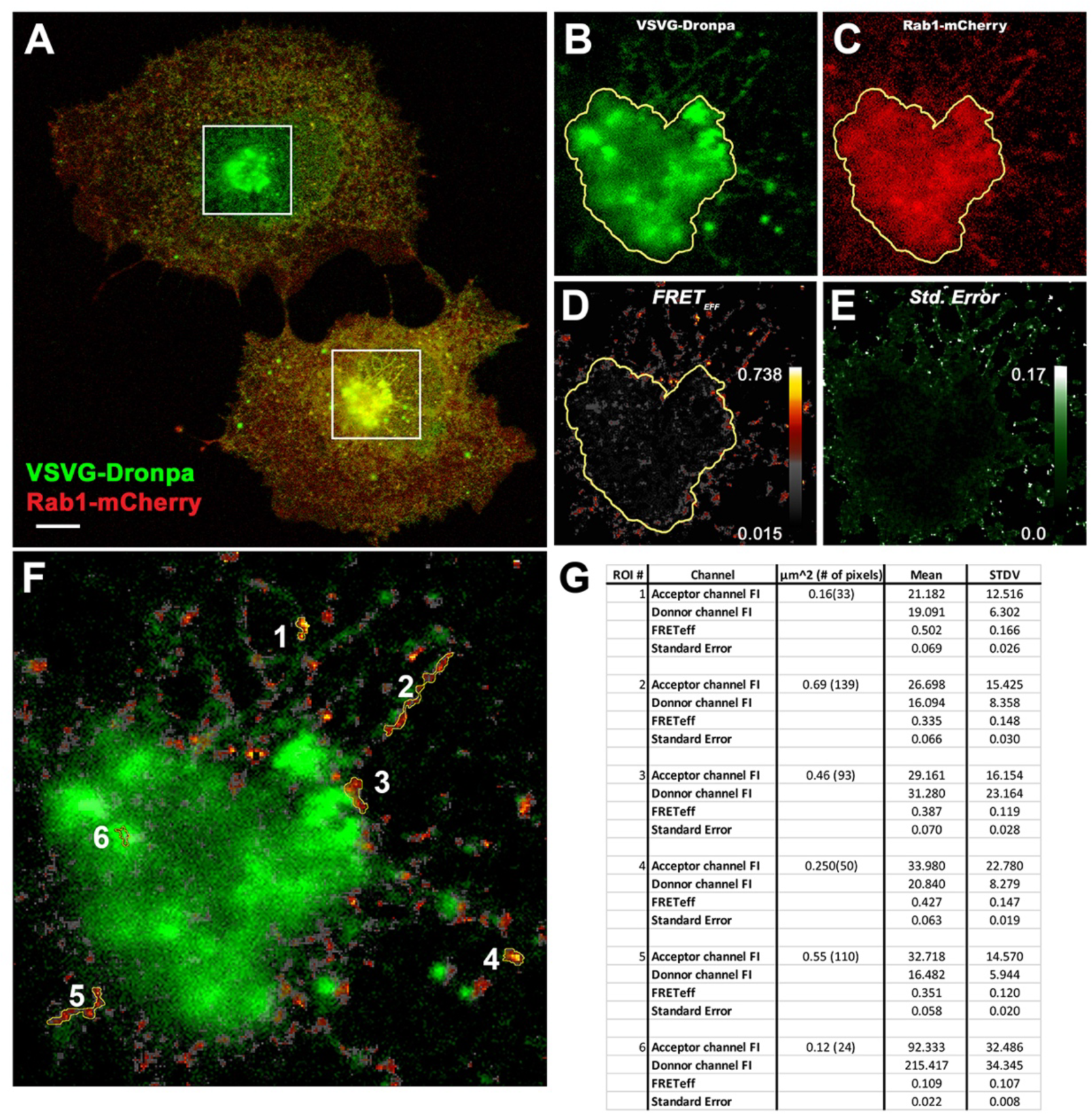
Rab1b and VSVG do not interact in Golgi membranes. COS7 cells co-expressing Rab1b-mCherry and VSVG-Dronpa were fixed after an incubation of 25 minutes at the permissive temperature of 32°C following overnight at the non-permissive temperature of 39.5 °C. **(A)** Confocal image of COS7 cells co-expressing VSVG-Dronpa (green) and Rab1b-mCherry (red). Top cell contains the bleached ROI used for determining the control exponential decay coefficient. bar = 10 µm **(B)** VSVG-Dronpa in the Golgi region enlarged 3-fold. **(C)** Rab1b-mCherry in the Golgi region enlarged 3-fold. **(D)** FRET efficiency interaction map in the Golgi region enlarged 3-fold. **(E)** Standard error of the fit in pseudo-color in the Golgi region enlarged 3-fold. **(F)** Analysis of FRET efficiency in ROIs associated with arriving carriers (1-5) and one within the Golgi (6). **(G)** A table depicting for each ROI in F, the values and standard deviations of donor and acceptor fluorescence, FRET efficiency, and standard error of the fit.

**Fig. 6.**
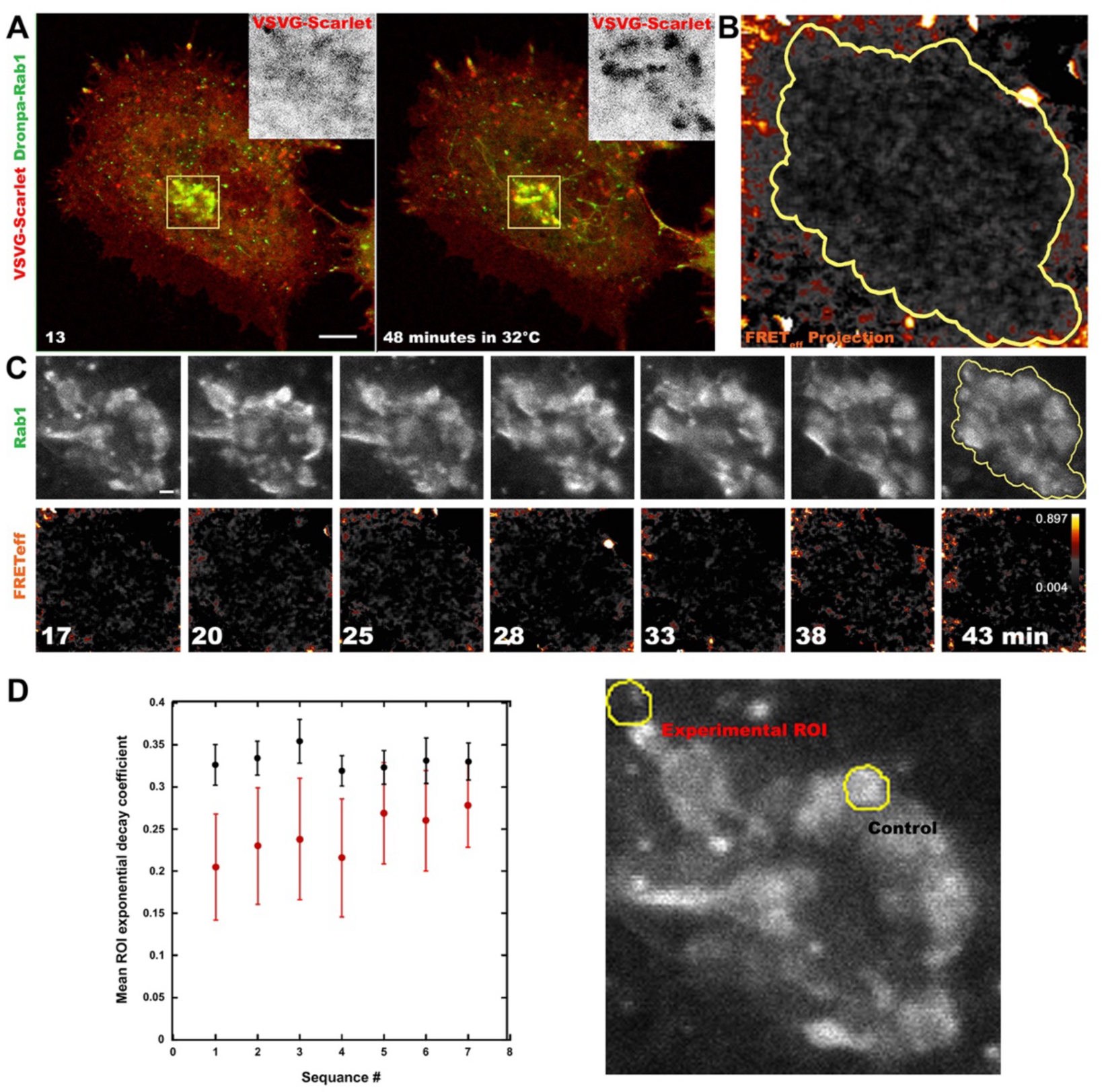
psFRET analysis of Rab1b-VSVG interaction in living cells. An ROI over the Golgi of COS7 cells co-expressing Rab1b-mCherry and VSVG-Dronpa was analyzed at seven timepoints starting from 17 to 43 minutes at the permissive temperature of 32°C following overnight at the non-permissive temperature of 39.5°C. **(A)** The cell co-expressing Rab1b-Dronpa (green) and VSVG-Scarlet (red) used for the psFRET analysis at 13 and 48 min after shift to the permissive temperature of 32°C. Yellow square is the ROI used. Top right insert is an inverted image of the VSVG-Scarlet channel. Scale bar = 10 µm. **(B)** Brightest pixel projection of seven consecutive live FRETeff interaction maps for the ROI over the Golgi of the cell in A. Yellow line delineates The Golgi apparatus. **(C)** Seven consecutive psFRET analyses obtained for the ROI within the living cell in A between the designated times after shift to permissive temperature. Top panel is the first image of the green Rab1b-Dronpa channel for each timepoint. Scale bar = 1 µm. Bottom panel is a pseudo color FRETeff interaction map. **(D)** A graph showing the exponential decay coefficient values ± standard error of the fit extracted from the two ROIs (yellow circles) at each timepoint. on the image at the right-hand side. Lower right ROI was used as the control values. Top left ROI included lower values signifying FRET.

### Rab1b interacts with the liquid-order domain promoting polytopic protein plasmolipin

VSVG has a single transmembrane domain and is considered a protein that favors a liquid-disordered membrane environment associated with basolateral membranes in epithelia (Ikonen and Simons, 1998). To test if Rab1b interacts with different cargo proteins, we analyzed Rab1b’s interaction with plasmolipin, a myelin and epithelial apical membrane protein with four transmembrane domains (Yaffe et al., 2015). Plas molipin belongs to the MARVEL protein family associated with, resides in, and promotes the formation of liquid-ordered membrane domains in the Golgi (Magal et al., 2009; Sanchez-Pulido et al., 2002; Yaffe et al., 2012). Supplementary Fig S4 demonstrates that Rab1b interacts with plasmolipin in pre-Golgi and Golgi membranes. Unlike VSVG, plasmolipin is a Golgi resident protein that recycles through the PM. Its interaction with Rab1b is surprising because plasmolipin resides and promotes the formation of liquid-ordered membrane domains. Based on the psFRET, the interaction in Golgi membranes is restricted to evenly distributed puncta.

### Live cell psFRET analysis

As the Dronpa can be switched on multiple times, we applied it to the analysis of psFRET in unfixed living intact cells coexpressing Rab1b-Dronpa and VSVG scarlet. The experiment was conducted at high confocal zoom to allow the fastest possible image capture times of about 20-50 milliseconds (Figure 6A). Fast image capturing is obligatory to capture the Dronpa decay image set while minimizing movements within this timeframe. Here, seven Dronpa decay stacks were taken of a small ROI over the Golgi of the same cell between 17 and 43 min after shifting to permissive temperature. Figure 6B shows the brightest pixel projection of all seven psFRET interaction maps. The psFRET interaction is restricted to pre-Golgi structures and is absent throughout the sequence from Golgi membranes. The individual psFRET images are shown in Figure 6C. To calculate the psFRET efficiency, we extracted the control decay value from an ROI over the Golgi membranes containing bright donor and acceptor fluorescence at all time points but with little or no FRET (Fig. 6D image on the right-hand side). The mean exponential decay coefficient values are shown for the experimental and control ROIs throughout the seven time points with standard error bars. Compared to the control ROI, the low exponential decay values indicate FRET interaction throughout the sequence.

### Overexpression of Rab1b inhibits Brefeldin A-mediated Golgi collapse into the ER

The context of cargo binding by Rab1 during ER to Golgi transport is unclear. Our hypothesis is that cargo interaction may increase the dwell time of the membrane associated active GTP-bound Rab1b resulting in increased recruitment of downstream effectors. To this end we tested the effect of overexpressed Rab1b-mCherry on BFA-mediated Golgi redistribution into the ER. Overexpression of Rab1b-mCherry increases the total levels of expressed Rab1b in the cells thereby increasing both cytosolic GDP-bound as well as the membrane bound GTP-bound active Rab1b. The latter may cause an increase in downstream effector recruitment. The COPI protein coat is recruited by the ARF1 GTPase for which GBF1, its GEF, is a downstream effector of Rab1b. To this end elevated membrane bound GTP-bound active Rab1b is expected to counter the effect of BFA-mediated release of ARF1 and COPI (Presley et al., 2002) leading to Golgi collapse. This effect is demonstrated in Fig. 7 showing that Golgi dispersion occurred in a cell that did not express Rab1b-mCherry compared to neighboring Rab1b-mCherry expressing cells where the Golgi was not affected. These data are in line with the hypothesis that Rab1b interaction with cargo may affect the level of downstream effectors recruitment. Thus, we introduced a comprehensive protocol, including a powerful analysis tool to perform psFRET. We applied the psFRET method to study fundamental interactions among components of the early secretory machinery (Fig. 8). Primarily, we localized Rab1b interaction with its downstream effector GRASP65. Remarkably, for the first time to our knowledge, direct interaction of Rab1b with the VSVG cargo protein was observed commencing after ER export and ending upon arrival at the Golgi apparatus.

**Fig 7.**
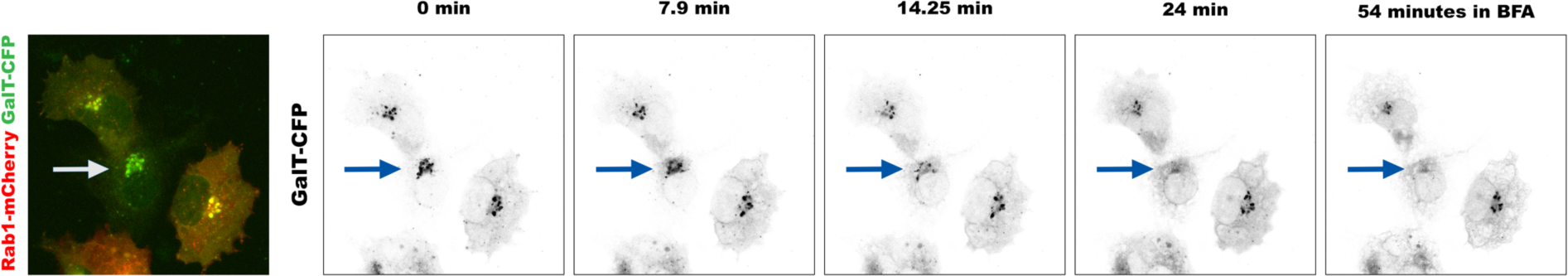
Rab1b overexpression prevents BFA-mediated Golgi redistribution into the ER. Living COS7 cells were co-transfected with Rab1b-mCherry (red) and GalT-CFP (green). Images of a field of view containing three cells with varying expression levels of Rab1b-mCherry were obtained using a confocal microscope (left merged image). Cells were treated with Brefeldin A and imaged at indicated time points following treatment (right GalT-CFP inverted images). Arrow indicates a cell without Rab1b overexpression.

**Fig. 8.**
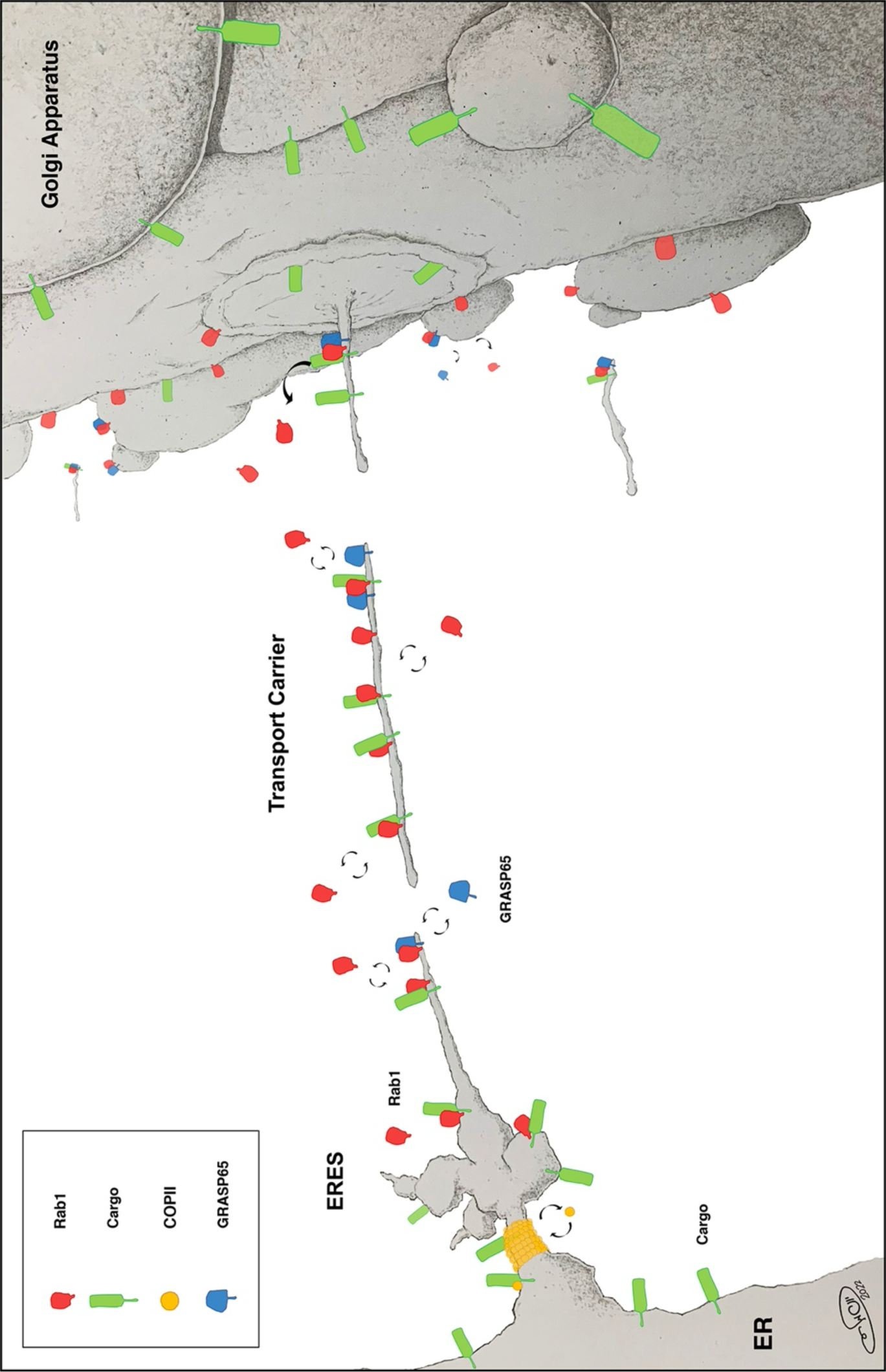
Schematic Representation of Rab1 interactions with cargo and downstream effectors during ER-to-Golgi transport. Newly synthesized cargo proteins in the ER are recognized by the COPII coat machinery at the ER-ERES boundary. Transport competent cargo proteins concentrate in the expending ERES while interacting with GTP-bound Rab1. GRASP65, a downstream effector of Rab1 is also recruited to distinct points at the ERES membranes. ERES membranes undergo fission to become anterograde transport carriers that translocate to fuse with the cis side of the Golgi. Rab1-cargo interaction is severed upon their entry to the Golgi apparatus.

## Discussion

### The psFRET method

We introduce a detailed protocol and a powerful analysis tool for the Photoswitching FRET method. This method has several noteworthy advantages over other approaches applied to detecting FRET. The first is that the FRET efficiency data is derived from the exponential decay constants of the Dronpa fluorescent protein and not from the fluorescence intensity data of the donor and acceptor as in acceptor photobleaching. In psFRET, the impact of fluorescence intensity is limited to the statistical parameters of the fit. The higher the intensity, the better the fit, which is essential when designing the experiment. To this end, the control decay coefficient used for calculating FRET efficiency should be derived from a high fluorescence intensity ROI. Another advantage of the psFRET is that the FRET data is calculated for each pixel separately, yielding a full image of the FRET efficiency by simply converting the output worksheet to a pseudo-color image using ImageJ (Schindelin et al., 2012). The software tool we introduce includes various image preprocessing options, including binning (flexible kernel size) and pooled or sliding kernels. Binning allows combining the data from surrounding pixels to improve data quality, although at the expense of resolution. For example, we used an 8 x 8 kernel to generate the control decay coefficient from 32 pixels simultaneously. This option allows for obtaining an optimized value for the control at the expense of the data resolution, which is not essential for the ROI used as a control. Supplementary Table S1 demonstrates the effects of combinations of two threshold values, three kernel sizes, and four image processing options applied on the same ROI on exponential decay coefficient value ± standard deviation. For example, we were using a 40,000 grey level threshold with averaging of a “sliding” 8 × 8 kernel, resulting in a decay coefficient value of 0.5032 ± 0.007 (56,352 pixels used), using a 20,000 grey level threshold with the maximum value option (instead of averaging) within a 2 x 2 kernel results in a decay coefficient value of 0.4850 ± 0.057 (16,384 pixels used).

In general, for the FRET_eff_ data used to generate interaction maps, we applied a sliding 2 x 2 kernel. Another key advantage of the psFRET is derived from the fact that the Dronpa is a photo-switchable protein that can be switched on multiple times using a 405 nm laser line, which is a standard in most commercial confocal microscopes. Thus, experiments can be repeated numerous times. Moreover, as demonstrated in Figure 6, psFRET analysis can be performed on living cells. However, currently, there are several concerns about performing psFRET on living cells. The first is the requirement to capture the input image stack fast enough to minimize movements that negatively affect the fitting process. To this end, faster image acquisition and larger kernels should be used. However, reduced intensity and larger kernels will affect data accuracy and resolution. Obtaining the control values may require a different approach than the one suggested for fixed cells. One should select the largest decay coefficients extracted from pixels with the highest fluorescence intensities of the Dronpa donor. A point to consider when applying the psFRET method is that threshold values should be used at an intensity value that will prevent fitting data in the background and pixels with particularly weak intensity. These cannot be fitted with sufficient accuracy. A high threshold may also prevent the software tool from extending the analysis time due to unnecessary calculations. The threshold can be determined using trial and error guided by the output’s statistical parameters.

### Rab1b interaction with the VSVG cargo

Using psFRET, we found a novel interaction between the Rab1b GTPase and the VSVG cargo protein. Particularly interesting is that the ER, on one side, and the Golgi apparatus, on the other, form clear borders that confine this interaction. Rab1b-VSVG interaction is detected upon exiting from the ER, proceeds during ER to Golgi transport, and is severed upon Golgi entry. This sharp spatiotemporal confinement was verified using reciprocal donor and acceptor FRET experiments and live cell analysis. In the case of plasmolipin, according to the psFRET analysis, the interaction continued within Golgi membranes but was restricted to distinct punctate structures. However, unlike VSVG which passes through the Golgi, plasmolipin is a Golgi resident protein. When overexpressed, it affects Golgi membrane composition to the point of blocking transport of membrane mismatched cargo (12). The interaction of Rab1b with cargo may serve as a stabilizing component to cargo-loaded membranes directly or via recruitment of downstream effectors such as the COPI heterocomplex (Kawamoto et al., 2002; Monetta et al., 2007). Moreover, this Rab1b-cargo interaction may act to stabilize the active GTP-bound membrane-associated form of Rab1b and, in so doing, affect the recruitment level of downstream effectors. A key downstream effector of Rab1b is GBF1, a GEF for ARF1. Thus, among its many functions, Rab1b may regulate and mediate interactions leading to transition from COPII-associated cargo at the ER-ERES collar to cargo-COPI at the ERES or ERGIC membranes. The nature of the Rab1b-cargo interaction is unclear. A third protein may be involved that still allows the proximity range of a FRET signal. This third protein may be excluded from the Golgi, thus causing Rab1b-VSVG dissociation, as seen by the apparent lack of FRET in Golgi membranes loaded with both Rab1b and VSVG. Change in membrane lipid composition may also facilitate the apparent Rab1b-VSVG dissociation upon arrival at the Golgi. The PM-targeted VSVG may partition to PM microdomains on Golgi membranes, while the GTP-bound Rab1b is associated with thinner, more disordered membranes. The COPI complex is recruited to ER to Golgi transport carriers. How early this occurs is still a debate (6); some claim it is on ERGIC, and some claim it is as early as ERESs. The details of Rab1b function are important as it is not only a master regulator of secretory transport but is involved in many pathogenic processes from viral infections (17-20) to genetic and neurodegenerative diseases.

## Materials and Methods

### Reagents and constructs

Unless otherwise stated, all reagents were purchased from Sigma Chemical Co. (St Louis, MO). Human Sec24B was subcloned into pmCherry-C1 (Clontech) using *Sal*I and *Bgl*II restriction sites and verified by sequencing. Human Rab1b-Dronpa and VSVG-Dronpa were provided by Dr George Patterson (NIH, Bethesda, MD). VSVG-Scarlet (Dukhovny et al., 2009) and DiHCRed-GRASP65 (Nevo-Yassaf et al., 2012) are described elsewhere.

### Cell Culture and transfection

COS-7 cells were grown at 37°C in a 5% CO_2_–humidified atmosphere. Cell cultures were maintained in Dulbecco’s modified Eagle’s medium (DMEM) supplemented with 10% (v/v) fetal bovine serum and penicillin and streptomycin (Biological Industries, Bet-Haemek, Israel). A final concentration of 1% (vol/vol) nonessential amino acids was added to the COS7 cells culture media. Polyethyleneimine ‘MAX’ transfection reagent (Warrington, PA, USA) was used following the manufacturer’s protocols for plasmid DNA transfections of sub-confluent COS7 cells. Confocal laser scanning microscopy experiments were conducted 18 to 24 h after transfection.

### Microscopy (fixed cells)

18 to 24 hours following transfection COS7 cells were fixed by the addition of formaldehyde to the medium to a final concentration of 4% and incubation for 10 min at room temperature. Cells were then washed twice with PBS and were imaged using FluoroBrite™ DMEM imaging media (ThermoFisher Scientific, Waltham, Ma).

### Live-cell microscopy

Cells were imaged in DMEM without phenol red but with supplements, including 20 mM HEPES, pH 7.4. Transfection and imaging were performed in a 35-mm glass-bottomed microwell dish (MatTek, Ashland, MA) or glass coverslips. A Zeiss LSM710 or LSM800 confocal laser-scanning microscope (Carl Zeiss MicroImaging, Jena, Germany). Fluorescence emissions resulting from 405-nm excitation for CFP, 488-nm excitation for GFP, and 543-nm excitation for mCherry were detected using filter sets supplied by the manufacturer. The confocal and time-lapse images were captured using a Plan-Apochromat ×63 1.4-numerical-aperture (NA) objective (Carl Zeiss MicroImaging). The temperature on the microscope stage was stable during time-lapse sessions using an electronic temperature-controlled airstream incubator. Images and movies were generated and analyzed using the Zeiss LSM Zen software and NIH Image and ImageJ software (W. Rasband, National Institutes of Health, Bethesda, MD).

### psFRET

#### Experimental protocol

The photoswitching FRET is based on the use of the Dronpa photoswitching protein. The FRET data is extracted by quantifying the decrease in the value of the exponential decay coefficient. This coefficient is obtained by fitting an exponential decay equation of the fluorescence intensity for each pixel in the image stack that records the decay of the Dronpa protein. Several key issues to consider will allow optimization of the FRET data. The quality of the fitting process depends on the level of the Dronpa fluorescence intensity. Thus, obtaining the decay data as a stack should begin when most, if not all, the Dronpa molecules are switched on. This state of the Dronpa can be achieved by scanning the field of view with a 405 nm laser line. In addition, to optimize the fit, a full dynamic range of pixel values should be used. Furthermore, most of the Dronpa molecules in the last image of the stack should be in their dark state. Below is a step-by-step protocol for performing a psFRET experiment with a Dronpa and red fluorescent proteins tagged donor and acceptor (Figure 1).

1. Locate a field of view containing two adjacent cells expressing adequate levels of Dronpa-tagged donor and red fluorescent protein-tagged protein as the acceptor. Preferably one with a neighbor cell with similar expression levels of the plasmid constructs. To avoid bleaching the Dronpa, use the red channel to locate the desired field of view.
2. Obtain a donor and acceptor channels image of the field of view to be used as an overlay for the FRET data presentation.
3. To determine the control values (without FRET) of the Dronpa decay coefficient, photobleach the red acceptor within a defined ROI using multiple scans with a high-power laser. The ROI should include the highest possible intensity of the donor. The higher the intensity, the more accurate the control value. A post-bleach image should be saved to allow the location of the bleached zone in the stack for analysis.
4. About 10 images should be taken using laser intensity, resulting in full conversion of the Dronpa to its dark state. Multiple stacks can be captured, providing that the Dronpa will be switched on using a 405 nm laser.

### Data analysis

#### The psFret utility

The psFRET software utility (Figure S1) is designed to allow the application of well-defined basic image processing steps before the psFRET analysis.

The input is in the form of any size tiff stack of 8 or 16-bit. There are two sets of image processing options: The first is a sliding filter that slides the selected kernel one pixel at a time generating an image made with the average, median, or Gaussian values. The second creates an image with pooled maximum, mean, or median values. Of note is that the first set outputs an image with the original pixel number, while the second set yields a smaller image with pixel numbers divided by the kernel size. Large kernels may yield more accurate fits, however, at the expense of the resolution. The processed images can be viewed before the analysis (Figure S1). Supplementary Table S1 demonstrates the result of several image processing options and kernel sizes on the exponential decay coefficient. A key step that mast to be determined is the threshold value below which the analysis will not be performed, and the value of 0 will be added instead as a default. The threshold values may be determined by error and trial so that pixels that are too low to allow accurate fit will not be analyzed. The importance of the threshold is derived from the fact that the output in the form of worksheets is converted to a pseudo-color image. Scattered values from inaccurate fits may cause misrepresentation of the data by the lookup table.

The utility output is in the form of four worksheets: The exponential decay coefficient generated by the fit, and the standard error, chi-square, and root mean square error (RMSE) value of the fit. All actions are recorded and saved in a file called “odel_log_MM-DD-YYYY_hhmmss.log”

### Converting worksheets to pseudo-color interaction maps and histograms

The ImageJ free software package is applied to convert worksheet text files to pseudo-color interaction maps as well as histograms. Briefly, worksheet saved as a text file is imported using the Import Text Image command. The Adjust brightness/contrast command is used to adjust the image dynamic range to a lookup table (Figure S3).

## Open code source

https://github.com/geekazaurus/fret_utilitypsFRETutility_v02. https://www.dropbox.com/s/xnm3v3bqd9t4pnp/psFRETutility_v02.exe?dl=0

## Acknowledgments

Many thanks to Ella Sklan, Department of Clinical Microbiology and Immunology, ^2^ Faculty of Medicine, Tel Aviv University for technical help. This work was supported by a grant from Fondation Jérôme Lejeune to KH. The work was also supported by a Tel Aviv University Aufzien Family Center for Prevention and Treatment of Parkinson’s Disease grant to KH.

