## supplemental figures and table for "Using Photoswitching FRET to Define the Interaction Boundaries between the Rab1b GTPase and Secretory Cargo"

### Supplementary figures

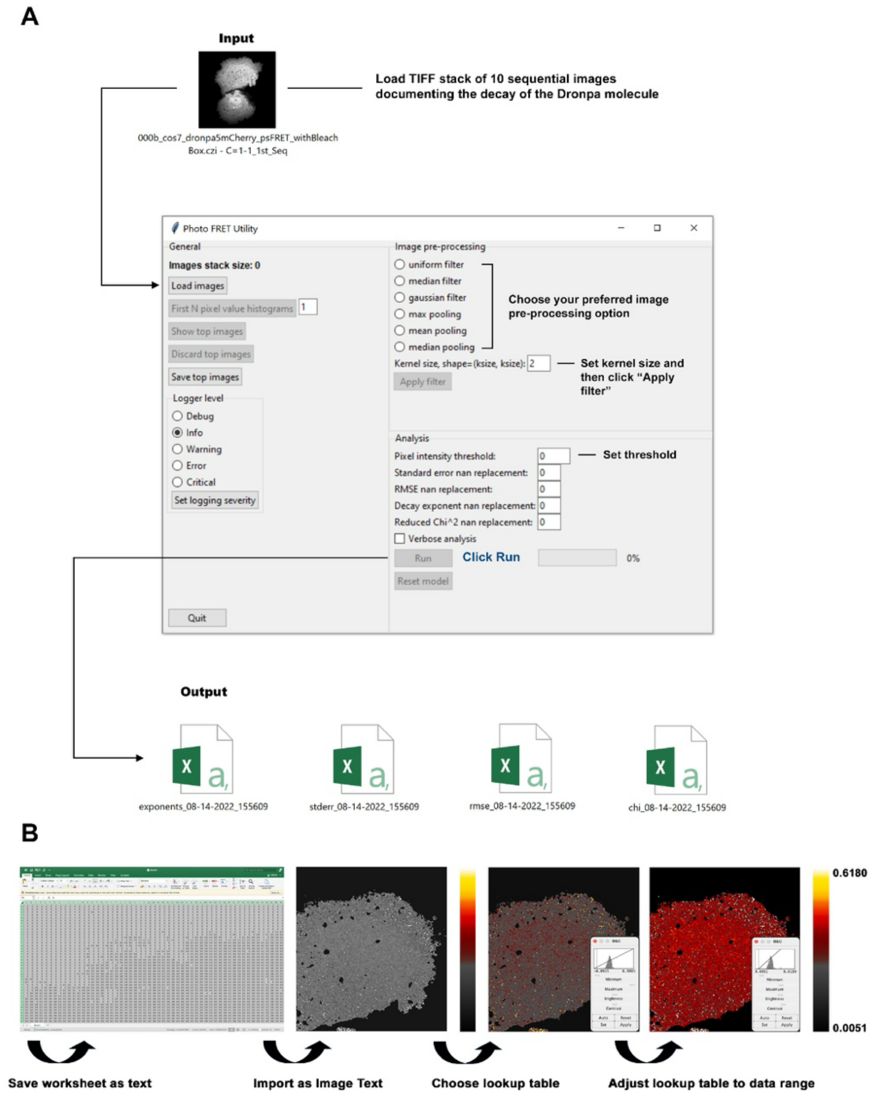


**Fig. S1. psFRET analysis and data presentation. (A)** The input to the analysis tool is in the form of a tiff stack. Preprocessing options are on the top right including fixed or sliding kernel size options. Effect of processing can be evaluated visually using "Show top images" button. Processed images can be discarded, and the input images can be processed repeatedly. Before the analysis a threshold value should be determined. Output is in the form of 4 worksheets one with the decay coefficients and three others with various error estimations. (**B)** Generation of interaction map from the psFRET utility using image J. The worksheet containing the exponential decay coefficients is saved as text after converting it to FRET efficiency values using Equation 1. The worksheet text file is opened using "Import Image Text" option. A lookup table can be selected and applied after which the Brightness & Contrast window is opened, and the dynamic range is adjusted to the positive range of the data.

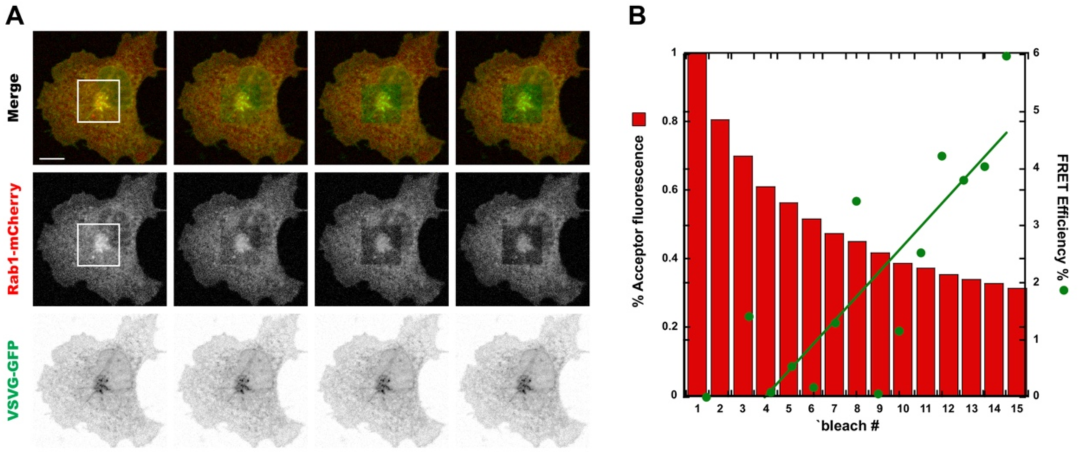


z

**Fig. S2. Stepwise acceptor photobleaching shows Rab1 and VSVG interaction. (A)** COS7 cell expressing Rab1-mCherry and VSVG-GFP was fixed using formaldehyde after an incubation of 25 minutes at the permissive temperature of 32°C following overnight at the non-permissive temperature of 39.5 °C. FRET was determined using stepwise acceptor photobleaching. Fluorescence of donor and acceptor were measured after each acceptor photobleaching. Intensities within the labeled (white square) region of interest were analyzed and are plotted in the graphs. Scale bar = 10 μm. **(B)** Quantification of the acceptor photobleaching FRET in A.

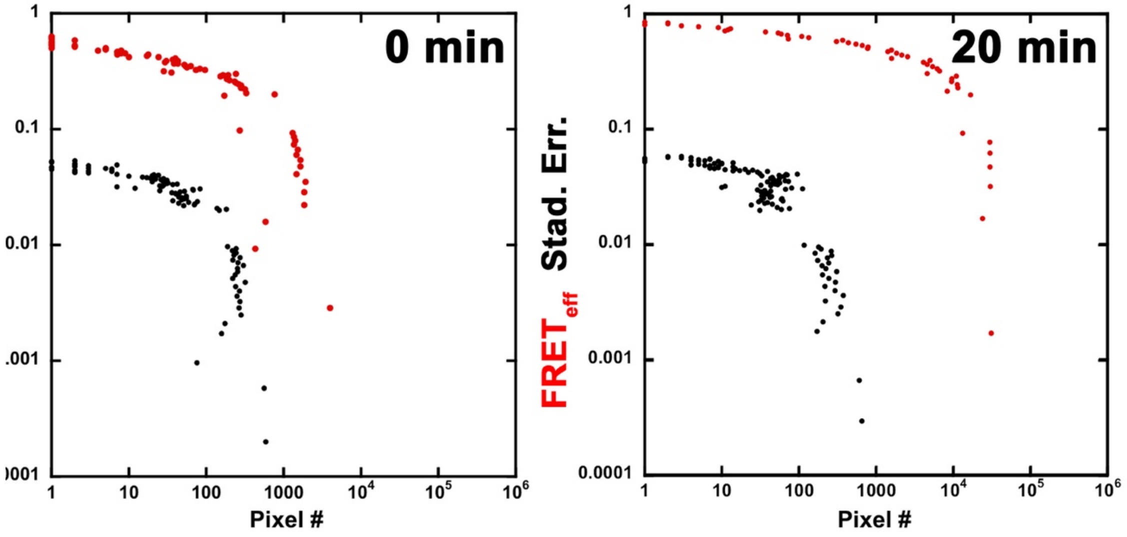


**Figure S3.** Log scale histogram analysis of Rab1b interaction with VSVG at 0 and 20 min after shift to permissive temperature (32 °C). Histograms show both FRETeff values (red filled dots) and standard error of the fit (black filled dots).

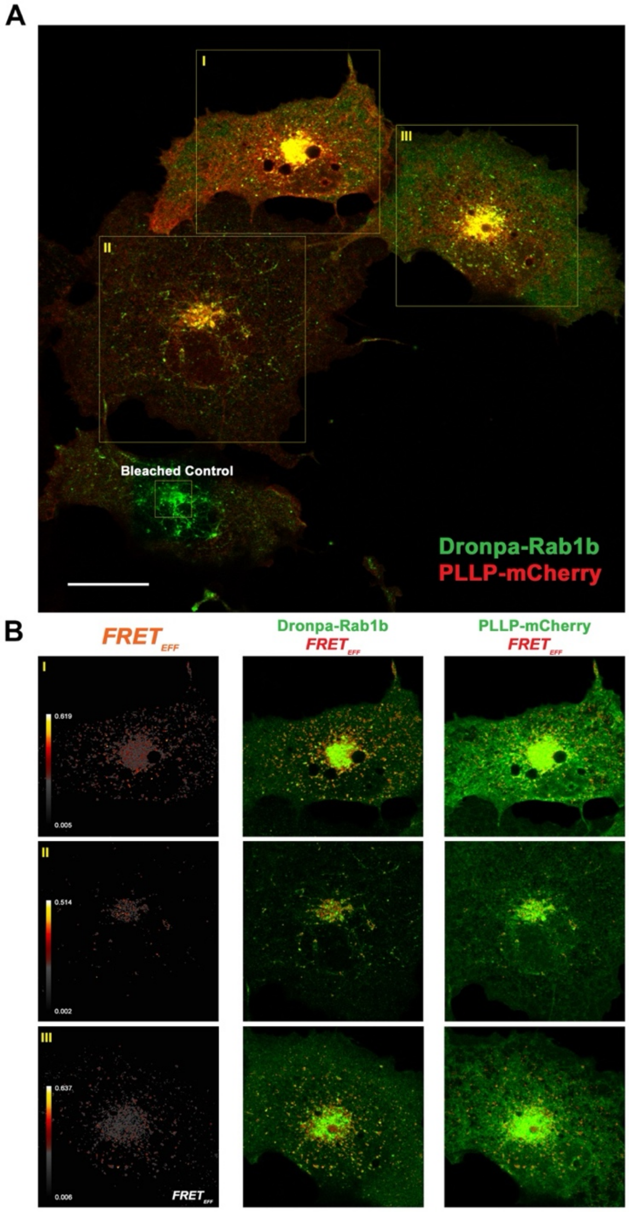


**Figure S4. Rab1b interacts with the liquid-order domain promoting polytopic protein plasmolipin. (A)** COS7 cells co-expressing Rab1b-Dronpa and PLLP-mCherry were fixed at steady state. Confocal image shows four cells and four ROIs one of which is the bleach-box (Bleached Control) over the Golgi apparatus of the cell at the bottom. ROIs I, II and III show three cells. **(B)** Left column is the FRET efficiency interaction map. Middle column is the interaction maps (red) overlayed on the Dronpa donor channel (green). Right column is the interaction maps (red) overlayed on the mCherry acceptor channel (green). Lookup tables show range of FRET efficiency. Scale bar = 20 μm.

| Uniform Filter - Threshold 20,000 | *k* (Mean) | Standard Deviation | Variance | Pixels used | | Uniform Filter - Threshold 40,000 | k (Mean) | Standard Deviation | Variance | Pixels used |
|---|---|---|---|---|---|---|---|---|---|---|
| **Kernel Size = 2** | 0.5021 | 0.024857631 | 0.0006179 | 65536 | | **Kernel Size = 2** | 0.5031 | 0.023642867 | 0.000559 | 54354 |
| **Kernel Size = 4** | 0.5015 | 0.012253917 | 0.0001502 | 65536 | | **Kernel Size = 4** | 0.5022 | 0.011441728 | 0.0001309 | 54761 |
| **Kernel Size = 8** | 0.5012 | 0.00806506 | 0.000065 | 65536 | | **Kernel Size = 8** | 0.5032 | 0.006846057 | 0.0000469 | 56352 |
| **Median Filter - Threshold 20,000** | **Mean** | **Standard Deviation** | **Variance** | **Pixels used** | | **Median Filter - Threshold 40,000** | **Mean** | **Standard Deviation** | **Variance** | **Pixels used** |
| **Kernel Size = 2** | 0.5013 | 0.029848276 | 0.0008909 | 65536 | | **Kernel Size = 2** | 0.5027 | 0.028527625 | 0.0008138 | 54855 |
| **Kernel Size = 4** | 0.5021 | 0.013490965 | 0.000182 | 65536 | | **Kernel Size = 4** | 0.5032 | 0.01258308 | 0.0001583 | 54598 |
| **Kernel Size = 8** | 0.5029 | 0.01003583 | 0.0001007 | 65536 | | **Kernel Size = 8** | 0.5052 | 0.008782658 | 0.0000771 | 55998 |
| **Gaussian Filter - Threshold 20,000** | **Mean** | **Standard Deviation** | **Variance** | **Pixels used** | | **Gaussian Filter - Threshold 40,000** | **Mean** | **Standard Deviation** | **Variance** | **Pixels used** |
| **Kernel Size = 2** | 0.5044 | 0.056317122 | 0.0031716 | 65536 | | **Kernel Size = 2** | 0.5085 | 0.053328147 | 0.0028439 | 53731 |
| **Kernel Size = 4** | 0.5017 | 0.015855301 | 0.0002514 | 65536 | | **Kernel Size = 4** | 0.5022 | 0.01507733 | 0.0002273 | 54660 |
| **Kernel Size = 8** | 0.5013 | 0.009419228 | 0.0000887 | 65536 | | **Kernel Size = 8** | 0.5028 | 0.008602967 | 0.000074 | 54689 |
| **Max Pooling - Threshold 20,000** | **Mean** | **Standard Deviaton** | **Variance** | **Pixels used** | | **Max Pooling - Threshold 40,000** | **Mean** | **Standard Deviation** | **Variance** | **Pixels used** |
| **Kernel Size = 2** | 0.485 | 0.057256833 | 0.0032783 | 16384 | | **Kernel Size = 2** | 0.4868 | 0.055544985 | 0.0030852 | 15015 |
| **Kernel Size = 4** | 0.4691 | 0.043659154 | 0.0019061 | 4096 | | **Kernel Size = 4** | 0.4694 | 0.042717757 | 0.0018248 | 3919 |
| **Kernel Size = 8** | 0.4549 | 0.038665093 | 0.001495 | 1024 | | **Kernel Size = 8** | 0.4546 | 0.038328674 | 0.0014691 | 1002 |
| **Mean Pooling - Threshold 20,000** | **Mean** | **Standard Deviation** | **Variance** | **Pixels used** | | **Mean Pooling - Threshold 40,000** | **Mean** | **Standard Deviation** | **Variance** | **Pixels used** |
| **Kernel Size = 2** | 0.5033 | 0.043885349 | 0.0019259 | 16384 | | **Kernel Size = 2** | 0.5059 | 0.041602638 | 0.0017308 | 13495 |
| **Kernel Size = 4** | 0.5021 | 0.024673264 | 0.0006088 | 4096 | | **Kernel Size = 4** | 0.503 | 0.023491162 | 0.0005518 | 3395 |
| **Kernel Size = 8** | 0.5017 | 0.016439778 | 0.0002703 | 1024 | | **Kernel Size = 8** | 0.5022 | 0.015774958 | 0.0002488 | 857 |
| **Median Pooling - Threshold 20,000** | **Mean** | **Standard Deviation** | **Variance** | **Pixels used** | | **Median Pooling - Threshold 40,000** | **Mean** | **Standard Deviation** | **Variance** | **Pixels used** |
| **Kernel Size = 2** | 0.504 | 0.048572706 | 0.0023593 | 16384 | | **Kernel Size = 2** | 0.5073 | 0.045913954 | 0.0021081 | 13457 |
| **Kernel Size = 4** | 0.5026 | 0.029241759 | 0.0008551 | 4096 | | **Kernel Size = 4** | 0.5042 | 0.027836356 | 0.0007749 | 3373 |
| **Kernel Size = 8** | 0.5023 | 0.018741895 | 0.0003513 | 1024 | | **Kernel Size = 8** | 0.503 | 0.01790685 | 0.0003207 | 850 |

**Table S1.** Effect of kernel size, preprocessing options, and threshold level on the control exponential decay coefficient in the same ROI.
